# Host ecological context influences taxonomic diversity and functional conservation of gut microbiome across anthropogenic habitats in macaques

**DOI:** 10.64898/2026.08.24.746638

**Authors:** Vani Kulkarni, Praveen Karanth, Sindhu Radhakrishna

**Author notes:** Corresponding author: Vani Kulkarni, Animal Behaviour and Cognition Programme, National Institute of Advanced Studies IISc Campus, Bengaluru 560012, India.

## Abstract

Gut microbiome responses to anthropogenic disturbance vary across wildlife species, even within similarly disturbed landscapes. What drives this variation is unclear: whether it reflects anthropogenic exposure itself or broader ecological differences among hosts. We tested this using three macaque species with contrasting ecology—Bonnet, Rhesus, and Lion-tailed macaques—sampled across 12 sites in southern India spanning contrasting anthropogenic exposure, using 16S rRNA gene sequencing (n = 127) and shotgun metagenomics on a subset of samples. The two synurbanized species exhibited a similar magnitude of microbiome restructuring but differed in the taxa underlying these changes; no differentially abundant amplicon sequence variants were shared across all three species, indicating that shared anthropogenic exposure did not produce uniform microbial responses across hosts. The specialist Lion-tailed macaque showed a more extensive response, characterized by reduced diversity and phylogenetically structured compositional change. The Bonnet macaque showed greater microbial similarity with the Rhesus macaque than with the Lion-tailed macaque during sympatric co-occurrence. Despite taxonomic divergence, functional pathway architecture was broadly conserved across species and habitats, with selective shifts in pathways including vitamin B6 biosynthesis and fermentation. Together, these findings show that microbiome responses to anthropogenic environments are jointly shaped by ecological context and host ecology, with host differences in diet, habitat use, and ecological history influencing the magnitude and nature of microbial restructuring. These findings show that taxonomic diversity and functional potential respond as partially decoupled axes under anthropogenic pressure, with implications for assessing microbiome resilience across ecologically heterogeneous wildlife.

## Introduction

Urbanization and anthropogenic ecological change are increasingly recognized as major forces reshaping wildlife biology, with host-associated gut microbiome emerging as a key interface through which these ecological drivers influence host physiology, health, and adaptive potential (Nguyen *et al*., 2024; Degregori *et al*., 2025; Peng *et al*., 2025). Gut microbiome plays an important role in host nutrition, metabolism, immune regulation, and adaptation to changing environmental and dietary conditions, thereby influencing the ecological flexibility and persistence of wildlife species under anthropogenic change (Ley *et al*., 2008; Barko *et al*., 2018; Clayton *et al*., 2018; Greene *et al*., 2020; Fackelmann *et al*., 2021). Across vertebrates, altered dietary ecology, habitat fragmentation, and intensified human–wildlife interactions can restructure gut microbiomes resulting in differential patterning of microbial taxa and function (Alberdi *et al*., 2016; Clayton *et al*., 2016; Di Rienzi and Britton, 2020; Moustafa *et al*., 2021; Nguyen, Lara-Gutiérrez and Stocker, 2021; Wasimuddin *et al*., 2022).

Microbiome restructuring can produce “taxonomic and/or functional microbial responses,” across related host species and among species occupying comparable ecological habitats, and/or, in some cases, reduced inter-host microbial distinctiveness within species (Yildirim *et al*., 2010; Clayton *et al*., 2016; Requena, Martínez-Cuesta and Peláez, 2018; Wasimuddin *et al*., 2022; Nguyen *et al*., 2024). Microbial alterations can also result in host-constrained restructuring, in which microbial communities retain host-associated taxonomic and functional composition despite comparable anthropogenic pressures. Such reshaping arises when host-associated ecological characteristics (e.g., genetics, immune system, and physiology) limit the extent of microbial patterning in response to external ecological pressures (Ley *et al*., 2008; Amato *et al*., 2016; Mazel *et al*., 2018; Somers *et al*., 2023; Marsh, Bearhop and Harrison, 2024). These restructuring processes are often scale-dependent, with microbial alterations varying across individuals, host systems, and ecological habitats (Adair and Douglas, 2017; McDonald, Marchesi and Koskella, 2020; Somers *et al*., 2023). Additionally, taxonomic and functional responses may not always occur in parallel, such that substantial compositional turnover can occur even when broader functional potential remains comparatively conserved due to microbial functional redundancy (Moya and Ferrer, 2016; Louca *et al*., 2018). Microbial responses may also manifest unevenly across abundance structure, phylogenetic membership, rare microbial taxa, and functional pathways, highlighting the multidimensional nature of these responses (Xia *et al*., 2022; Somers *et al*., 2023; Ma, 2026).

Anthropogenic exposure is a prominent driver of gut microbiome restructuring in wildlife, but microbial responses vary considerably across vertebrate taxa and ecological contexts (Littleford-Colquhoun *et al*., 2019; Maraci *et al*., 2022; Adair *et al*., 2025; Peng *et al*., 2025). For example, urban-associated reptiles and some birds exhibit increased microbiome diversity, whereas some avian and mammalian hosts more commonly show reduced diversity, compositional restructuring, or enrichment of pathogenic taxa under human-modified conditions (Sugden, St. Clair and Stein, 2021; Dillard *et al*., 2022; Tsuchida *et al*., 2023; Adair *et al*., 2025). In primates, responses range from reduced alpha diversity and depletion of fiber-degrading taxa to compositional reshaping without measurable diversity change (Lee *et al*., 2019; Chen *et al*., 2020; Wasimuddin *et al*., 2022; Liu *et al*., 2024). This variability likely reflects the interplay between habitat-level ecological pressures and host-specific ecological histories, yet existing studies have been poorly positioned to disentangle them (Somers *et al*., 2023; Liu *et al*., 2024; Degregori *et al*., 2025; Peng *et al*., 2025). Most investigations have relied on single-species frameworks with single-axis habitat contrasts — urban versus wild, provisioned versus non-provisioned, that combine environmental signals with host-specific responses, leaving it unclear whether observed microbiome shifts reflect shared anthropogenic pressures or species-specific adaptations (Chen *et al*., 2020; Wasimuddin *et al*., 2022; Amato *et al*., 2025; Beeby, Pierre and Guy, 2025). Even comparative studies involving sympatric species have largely emphasized taxonomic restructuring, with functional pathway composition and the relationship between taxonomic and functional responses receiving comparatively limited attention (Moeller *et al*., 2013; Liu *et al*., 2021; Anders *et al*., 2022; Heni *et al*., 2023; Tuoliu *et al*., 2024). It therefore remains unclear whether comparable anthropogenic conditions promote similar taxonomic and functional microbiome responses in related species with differing ecological flexibility, or whether host ecology maintains distinct microbial structures even among sympatric species.

Among primates, macaques (genus: Macaca) exploit anthropogenic resources across gradients ranging from forest and forest-edge habitats to peri-urban and urban habitats, yet species differ substantially in ecological flexibility and extent of anthropogenic association. Bonnet macaques (*Macaca radiata,* BM hereafter) and Rhesus macaques (*Macaca mulatta,* RM hereafter) are synurbanized species exhibiting high behavioural flexibility and extensive reliance on anthropogenic food resources, though they differ in the nature of this integration: RM are characterized by broad geographic expansion and strong urban dominance, whereas BM persist primarily within fragmented peri-urban and human-modified mosaics in peninsular India (Kumar et al. 2011; A Sengupta and Radhakrishna 2016; Erinjery et al. 2017; A. Sengupta et al. 2025; Radhakrishna et al. 2025). Lion-tailed macaques (*Macaca silenus*; LTM hereafter) seemingly represent an ecological contrast, they are rainforest specialists with a restricted geographic range and strong sensitivity to habitat disturbance. However observations from fragmented habitats indicate emergent flexibility in anthropogenic resource use (Singh, 2019; Dhawale, Kumar and Sinha, 2020; Mahato *et al*., 2026).

A comparative framework involving these three macaque species provides a unique opportunity to examine gut microbiome structuring at multiple levels (individual, host-species, and interspecific within-genus) while minimizing phylogenetic variation. Comparing two synurbanized macaque species occupying similar anthropogenic environments allows assessment of whether anthropogenisation drives similar microbial responses or whether species-specific characteristics constrain microbial composition. Studying microbial responses in the rainforest-specialist macaque enables examination of how anthropogenisation modifies microbial patterning in an ecological specialist species. Using this framework, we investigated: (1) does anthropogenic exposure promote microbial similarity in macaques that differ in ecological flexibility, or does host ecology generate species-specific restructuring patterns?, (2) does sympatric co-occurrence promote microbiome similarity among macaque species with different degrees of ecological flexibility?, (3) does microbial taxonomic diversity and functional composition respond in a similar manner to anthropogenic influences?. We predicted that (1) the two synurbanized macaque species would exhibit a broadly similar magnitude of microbiome restructuring in response to anthropogenic exposure, whereas the habitat-specialist lion-tailed macaque would exhibit a more extensive, distinct response, (2) sympatric macaque species sharing anthropogenic habitats would exhibit similarity in microbial composition, and (3) microbial response patterns in terms of functional structure would be relatively more conserved than taxonomic composition.

## Methods

The study was conducted across 12 study sites in southern India between March 2024 and March 2025 (Fig. S1). The research protocol was approved by the Institutional research ethics committee of the National Institute of Advanced Studies. Necessary research permits were obtained from the state forest departments to conduct fieldwork and collect fecal samples from the three macaque species. The BM is endemic to southern India and is an adaptable generalist omnivore commonly inhabiting human-dominated landscapes such as temples, villages, agricultural areas, and urban settlements. The RM is widely distributed across northern India and is an opportunistic omnivore known for ecological flexibility and extensive use of anthropogenic environments. The LTM is a rainforest specialist endemic to the Western Ghats in southern India and is primarily associated with tropical evergreen forests (Singh, 2019). Sampling sites were selected to represent a range of environmental gradients from protected rainforest and deciduous forest habitats, to anthropogenic disturbed forest sites, rural regions, temple sites, and human-dominated tourist sites (Fig. S1). The sampled populations differed in the extent of human contact, access to anthropogenic food resources, provisioning opportunities, and reliance on natural versus human-derived resources. Two study sites supported sympatric occurrence of two macaque species, with one site (H1) comprising BM and LTM and another (H7) comprising BM and RM (Table S1).

### Fecal sample collection

Field observations were conducted approximately 10–12 days at each site, typically across consecutive sampling days. At each site, troops were initially located by walking pre-identified access routes and forest trails, often with the assistance of local field guides familiar with troop ranging patterns. Once located, troops were followed through the day during observation periods, allowing fresh fecal samples to be assigned to identified individuals/social groups after defecation. Across study sites, BM troop sizes ranged from 15–40 individuals, RM troops ranged from 50–150 individuals, and LTM groups from 15–50 individuals. In BM and LTM groups, moderate group sizes allowed reliable tracking of individuals during sampling, permitting collection of one sample per individual. RM groups were substantially larger and field conditions limited consistent tracking of individuals across days. Consequently, repeated sampling of the same individual across days in these groups could not be completely excluded. Fresh fecal samples were collected from the ground immediately after defecation while ensuring that samples had not been contaminated by soil or vegetation. A total of 127 samples were collected across three macaque species (BM = 45, RM = 46 and LTM = 36). To minimize environmental contamination, the outer surface of the fecal bolus was gently opened and the interior portion was swabbed using sterile swabs. Samples were preserved in a nucleic acid stabilization buffer and stored at −20°C until further processing.

### Microbiome DNA Extraction and Sequencing

Microbial DNA was extracted using the Qiagen QIAamp PowerFecal Pro Kit following manufacturer instructions. DNA quantity and purity were assessed using a Nanodrop spectrophotometer. The V3–V4 region of the bacterial 16S rRNA gene was amplified using primers with Illumina adapter overhang sequences and heterogeneity spacers (Naik *et al*., 2023). Libraries were prepared using a two-step PCR protocol and sequenced on an Illumina MiSeq platform (2 × 300 bp) at the Next Generation Genomics Facility, NCBS, in Bengaluru. A total of 127 fecal samples were sequenced, which generated a total of 34,926,109 reads, with an average of 275,009 reads per sample prior to filtering.

Sequence data were processed using QIIME2 (version 2024.10) (Bolyen *et al*., 2019). Demultiplexed reads were quality filtered, denoised, and merged using the DADA2 plugin (trimming: 270/220 bp) (Callahan *et al*., 2016). Taxonomic classification was performed using the QIIME2 classify-sklearn plugin with the SILVA (v138) reference database. Amplicon sequence variants (ASVs) classified as mitochondria or chloroplast were removed. After filtering and chimera removal, the dataset contained an average of 44,992 sequences per sample (range: 1,043–81,269), resulting in 9,584 ASVs across the dataset. Samples were rarefied to 9,500 reads based on inspection of rarefaction curves to balance sequencing depth and sample retention. Phylogenetic trees were constructed using MAFFT within the QIIME2 pipeline.

### Anthropogenic Exposure Framework

We classified study sites as High Anthropogenic Exposure (HAE) and Low Anthropogenic Exposure (LAE) categories based on habitat type, nature and source of food resources and human–macaque interaction intensity. HAE sites (n = 7) were characterized by ecological contexts in which provisioning, food raiding, roadside feeding, or tourist- and temple-associated food access constituted a substantive component of the foraging environment. LAE sites (n = 5) were characterized by predominantly natural foraging contexts with minimal anthropogenic food input.

### Statistical analysis

All analyses were conducted in R (v4.5.2) (R Core Team, 2025). Alpha diversity (Observed ASV richness, Shannon index, Faith’s PD) was calculated using phyloseq (richness/Shannon) and picante (Faith’s PD), and compared between environmental categories within each host species using Wilcoxon rank-sum tests (McMurdie and Holmes, 2013).

Beta diversity (Bray–Curtis, weighted/unweighted UniFrac) was calculated from the rarefied ASV dataset and visualized via PCoA (ape package). Community composition differences were tested with PERMANOVA (adonis2, vegan, 9,999 permutations), modeling host species, habitat category (high vs. low anthropogenic exposure), and their interaction; multivariate dispersion homogeneity was checked with betadisper before interpreting results. The same beta diversity/PCoA/PERMANOVA/betadisper workflow was repeated on a subset of sympatric host-species samples (zero-abundance taxa removed post-subsetting). Interspecific Bray–Curtis dissimilarities were classified as sympatric or non-sympatric and compared overall and per species pair (BM–RM, BM–LTM) via Wilcoxon rank-sum tests; HAE–HAE vs. LAE–LAE interspecific distances were similarly compared to assess whether effects of co-occurrence are independent of habitat category. Interspecific distances were further contextualized against conspecific different-site distances using Kruskal–Wallis with Benjamini-Hochberg-corrected pairwise Wilcoxon tests.

Differential abundance was assessed at the ASV level with ANCOM-BC2 (Lin and Peddada, 2024) and ALDEx2 (Gloor, 2023), after removing non-bacterial/chloroplast/mitochondrial features and ASVs present in <10% of samples. ANCOM-BC2 (habitat category and host species as fixed effects) was run globally and separately within each macaque species (q < 0.05, BH-corrected). ALDEx2 (CLR transformation, Monte Carlo sampling, Welch’s t-tests) served as a compositional-robustness check. Differentially abundant ASVs were compared across species/methods via set intersections and collapsed to family level for taxonomic summary.

A representative 40-sample subset (BM = 10, RM = 15, and LTM = 15) spanning all three host species and both anthropogenic exposure categories underwent shotgun metagenomic sequencing to balance sequencing costs while retaining ecological representation. Following host-read removal, microbial functional profiling was performed using HUMAnN (v3.9), which first identified microbial taxa using MetaPhlAn (v4.2.4) and then mapped reads to the ChocoPhlAn pangenome database, with unmapped reads searched against the UniRef90 protein database using DIAMOND to reconstruct MetaCyc pathway abundances. Functional community structure was assessed using Bray–Curtis dissimilarity, PCoA, and PERMANOVA (vegan), and differential pathway abundance was tested using ANCOM-BC2.

## Results

### Anthropogenic exposure restructures gut microbial composition across macaque species

Across all macaque species and environmental categories, gut microbial communities were dominated by Lachnospiraceae, Prevotellaceae, Ruminococcaceae, Oscillospiraceae, Spirochaetaceae, and Rikenellaceae, with the remaining 99 bacterial families collectively accounting for 45% of the mean community composition (Fig. 1). BM and LTM showed similar microbial responses; in HAE habitats there were increases in Prevotellaceae (BM: 13.1% to 16.8%, LTM: 9.1% to 12.7%) and Spirochaetaceae (BM: 2.6% to 4.9%; LTM: 1.4% to 5.5%) and declines in Ruminococcaceae (BM: 11.6% to 8.7%, LTM: 9.7% to 7.3%) and Lachnospiraceae (BM: 17.3% to 14.4%, LTM: 16.0% to 14.1%). Rhesus macaques diverged from this pattern, showing a decline in Prevotellaceae under HAE conditions (20.3% to 15.8%), and increase in Oscillospiraceae (3.3% to 4.9%) and Rikenellaceae (1.4% to 3.2%).

**Figure 1.**
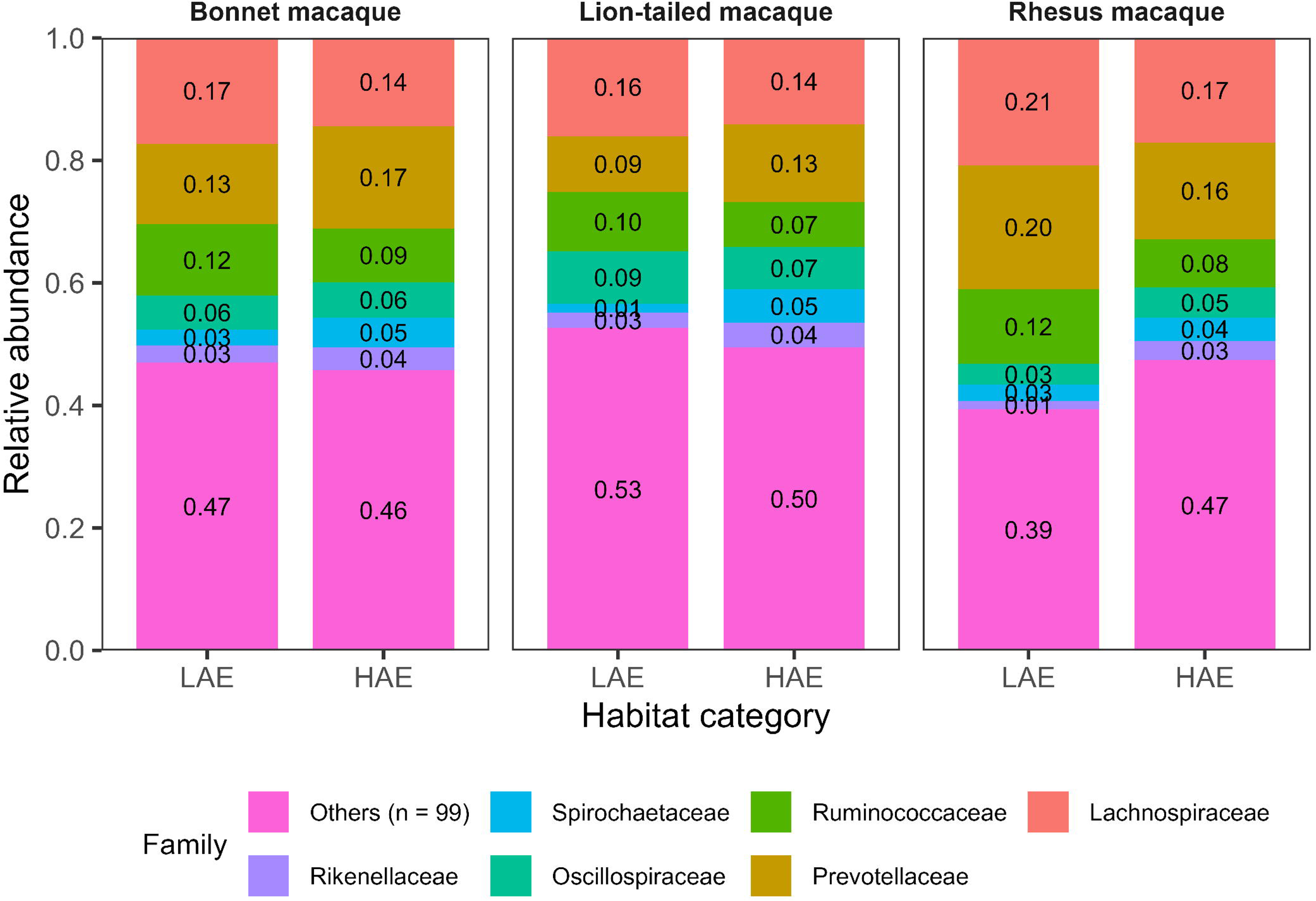
Relative abundance of the dominant bacterial families in bonnet macaques (BM), lion-tailed macaques (LTM), and rhesus macaques (RM) across low-anthropogenic-exposure (LAE) and high-anthropogenic-exposure (HAE) environments. Stacked bars represent the mean relative abundance of bacterial families within each host species and anthropogenic environment. “Others” represents the pooled contribution of the remaining 99 bacterial families.

Alpha diversity responses to anthropogenic exposure varied markedly among host species (Fig. 2). LTM exhibited significant differences across all alpha diversity metrics, including Observed ASV richness (p = 0.0006), Shannon diversity (p = 0.00032), and Faith’s phylogenetic diversity (p = 0.0019). Individuals from LAE environments exhibited higher microbial richness, diversity, and phylogenetic diversity than those from HAE environments, indicating reduced within-host microbial diversity in response to higher anthropogenic exposure. BM and RM showed no significant differences in alpha diversity across environmental categories (all p > 0.05) (Fig. 2).

**Figure 2.**
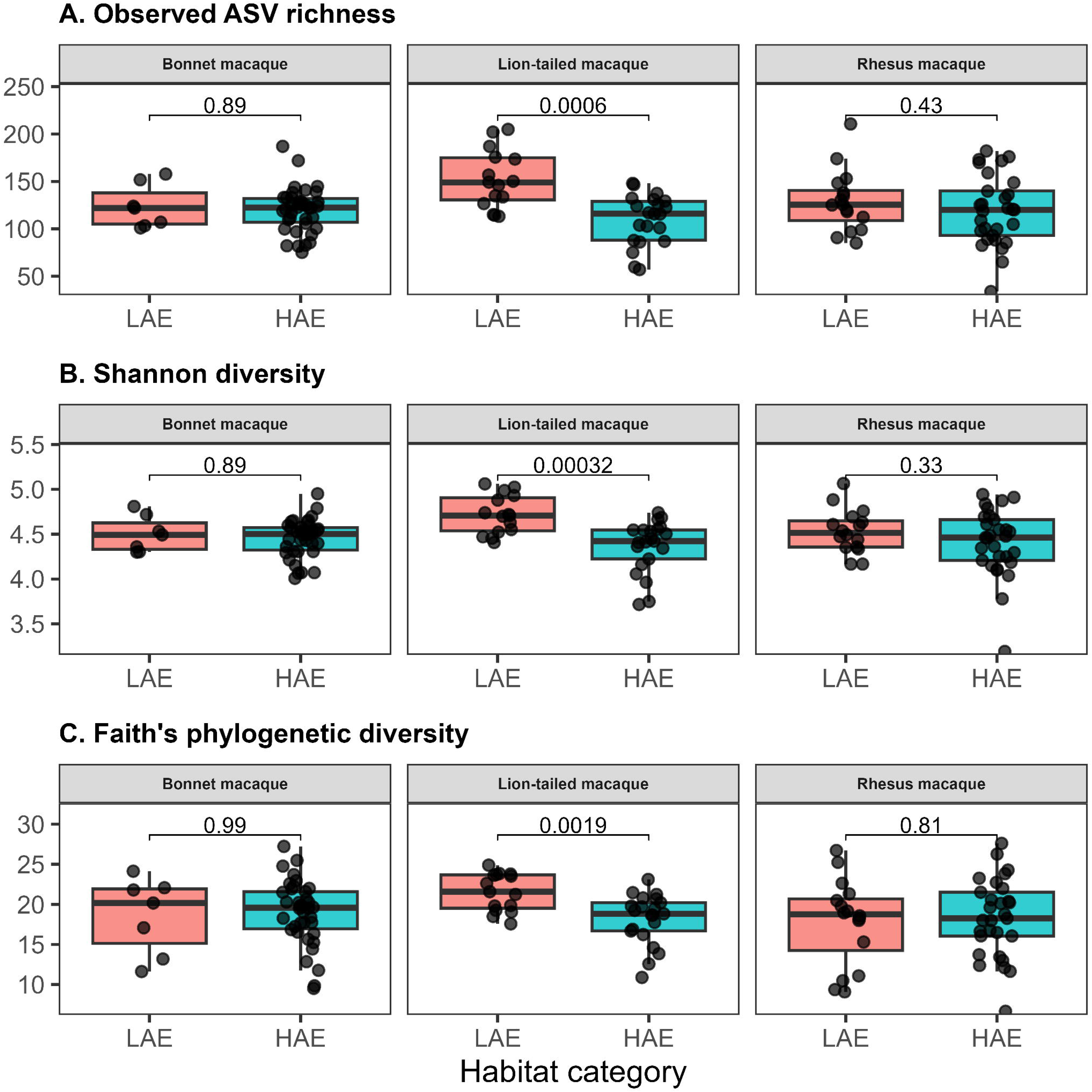
Alpha diversity responses across environmental categories. Observed ASV richness (A), Shannon diversity (B), and Faith’s phylogenetic diversity (C) in BM, LTM, and RM inhabiting low- and high-anthropogenic-exposure habitats. Points represent individual samples and boxplots indicate median and interquartile range.

Beta diversity analyses revealed that habitat-associated microbiome responses differed by host species. A global host × habitat interaction model confirmed this species-dependence (Bray– Curtis: R² = 0.024, p = 0.0001; unweighted UniFrac: R² = 0.025, p = 0.0069; weighted UniFrac: p = 0.324; Table S2, Fig. S2). The significant unweighted-but-not-weighted UniFrac result suggested that habitat-associated shifts were more evident among lower-abundance taxa than among dominant community members. Species-specific comparisons showed that BM and RM exhibited a similar magnitude of response, with significant habitat-associated differentiation across all three beta-diversity metrics. Both species also showed significant Bray–Curtis dispersion heterogeneity, indicating that habitat-associated compositional shifts were accompanied by greater within-habitat variability rather than a consistent directional change. LTM, by contrast, showed significant differentiation across all three metrics without dispersion heterogeneity, indicating a consistent shift in community composition across habitats (Table 1, Table S4; Fig. 3).

**Figure 3.**
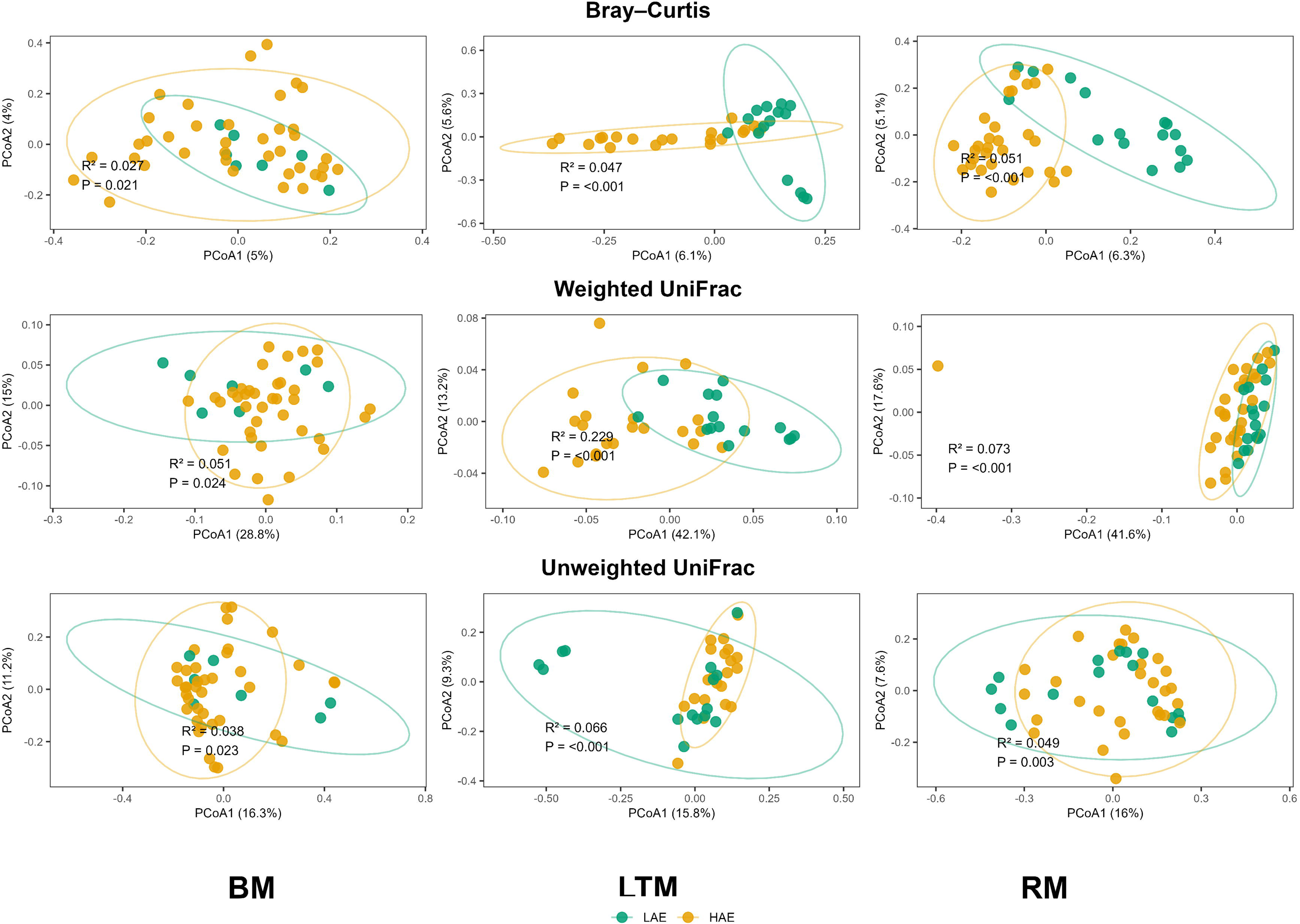
Principal coordinates analysis (PCoA) of gut microbiome beta diversity in BM, LTM, and RM across high (HAE) and low (LAE) anthropogenic exposure environments. Rows show ordinations based on Bray–Curtis dissimilarity (top), weighted UniFrac distance (middle), and unweighted UniFrac distance (bottom); columns show each host species. Points represent individual samples, colored by habitat category (LAE, teal; HAE, orange), with ellipses denoting 95% confidence intervals for each group. R² and p-values are from PERMANOVA testing for habitat-associated differences in community composition within each species and metric (see Table 1 for full statistics, including dispersion tests).

**Table 1.** Species-specific PERMANOVA and multivariate dispersion analyses of gut microbial beta diversity across low- and high-anthropogenic-exposure (LAE and HAE) environments in BM, RM, and LTM.

| Host Species | Metric | R<sup>2</sup> | p | Betadisper p-value |
| --- | --- | --- | --- | --- |
| BM | Bray-Curtis | 0.027 | 0.021 | 0.001 |
| BM | Weighted UniFrac | 0.051 | 0.024 | Non-Significant |
| BM | Unweighted UniFrac | 0.038 | 0.023 | Non-Significant |
| RM | Bray-Curtis | 0.051 | 0.001 | 0.001 |
| RM | Weighted UniFrac | 0.073 | 0.001 | Non-Significant |
| RM | Unweighted UniFrac | 0.049 | 0.003 | Non-Significant |
| LTM | Bray-Curtis | 0.047 | 0.001 | Non-Significant |
| LTM | Weighted UniFrac | 0.229 | 0.001 | Non-Significant |
| LTM | Unweighted UniFrac | 0.066 | 0.001 | Non-Significant |

### Microbial restructuring versus host-constrained microbial composition across habitat categories

Differential abundance analyses identified 27 globally significant ASVs across habitat categories, of which 16 were enriched in high-anthropogenic-exposure (HAE) sites and 11 were enriched in low-anthropogenic-exposure (LAE) sites (Table S3). Microbial shifts were primarily associated with ASVs belonging to the Lactobacillaceae, Prevotellaceae, Lachnospiraceae, and Ruminococcaceae families (Fig. 4A). HAE-associated changes were dominated by Lactobacillaceae (n = 6 ASVs; mean logFC = 0.50) and Prevotellaceae (n = 5 ASVs; mean logFC = 0.62), whereas LAE-associated changes were characterized predominantly by Lachnospiraceae (n = 4 ASVs; mean logFC = −0.66). ASVs assigned to Prevotellaceae and Ruminococcaceae were associated with both HAE and LAE environments, suggesting that habitat-associated responses occurred at a finer taxonomic resolution than the family level.

**Figure 4.**
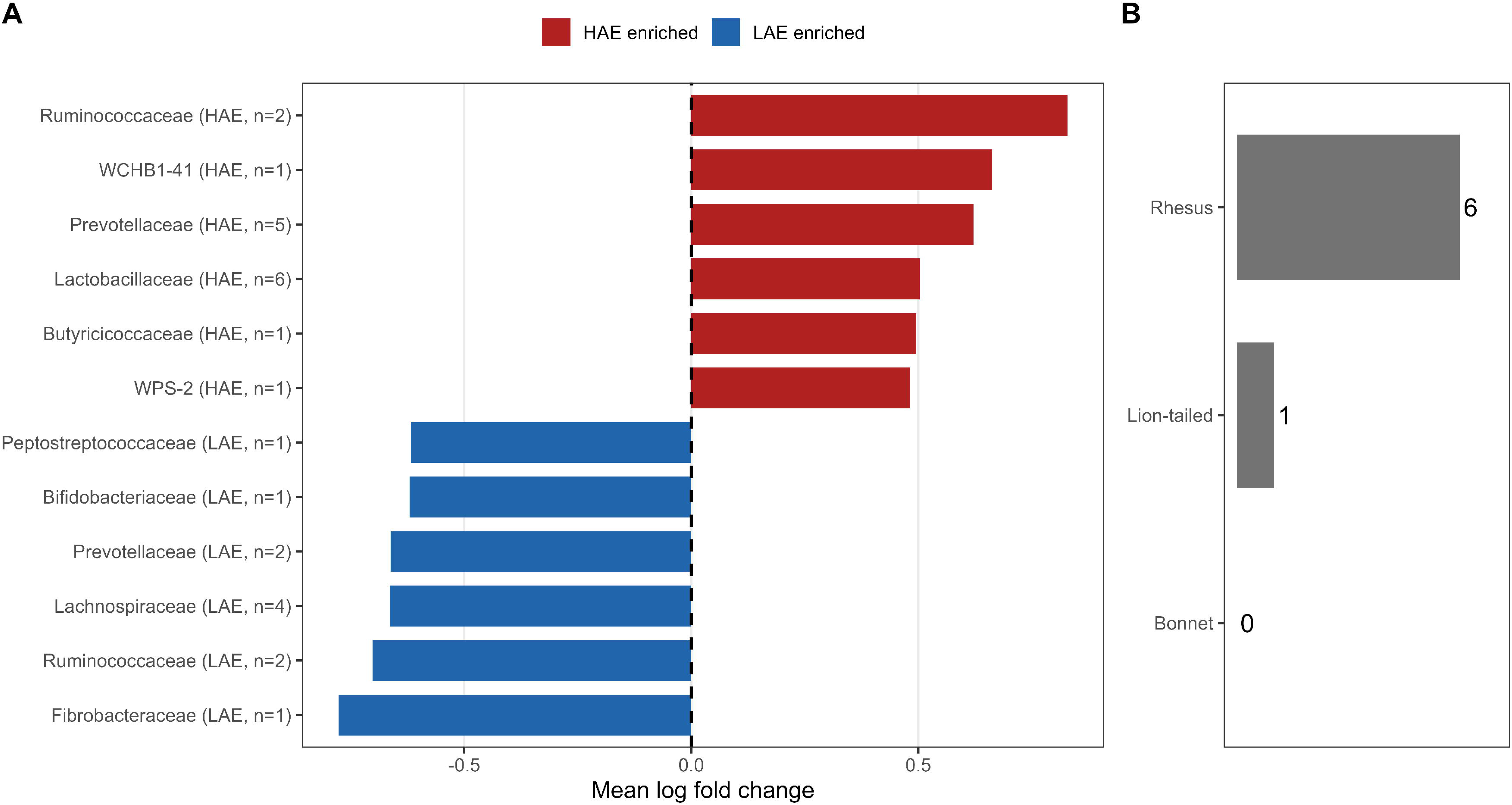
Differential abundance patterns across anthropogenic environments and macaque hosts. (A) Bacterial families significantly associated with high-anthropogenic-exposure (HAE) and low-anthropogenic-exposure (LAE) environments. Bars represent the mean log fold change of significant ASVs within each bacterial family; labels indicate the number of significant ASVs assigned to each family. (B) Number of significant differentially abundant ASVs identified within each macaque species.

Species-specific differential abundance analyses revealed no significant ASVs in BM, one significant ASV in LTM, and six significant ASVs in RM (Fig. 4B). No significant ASVs were shared between any pair of host species, and no single bacterial family contained significantly differentially abundant taxa in all three macaque species simultaneously, although individual families were sometimes represented among the significant taxa of one or two species. These patterns indicate that environmental influences did not produce extensive ASV-level overlap or diminish host-associated distinctiveness across these macaque microbiomes, despite exposure to broadly comparable ecological conditions.

Site-level comparisons assessed microbiome differentiation between sympatric macaque species (Fig. 5). In the H1 site, where BM and LTM co-occurred, there were significant differences between the host species with respect to Bray–Curtis (PERMANOVA: R² = 0.049, F = 1.61, p = 0.0002) and weighted UniFrac distances (R² = 0.065, F = 2.15, p = 0.022), but no significant differences in multivariate dispersion (Bray–Curtis: p = 0.745; weighted UniFrac: p = 0.652), indicating that host species differed in microbiome composition, and that these differences were not driven by differences in dispersion (Fig. 5A). In contrast, sympatric BM and RM in the H7 site did not differ in Bray–Curtis (p = 0.367) or weighted UniFrac distances (p = 0.583), but differed significantly in Bray–Curtis dispersion (p = 0.033), indicating greater within-group heterogeneity (Fig. 5B). Together, these results show that species-associated microbiome differences persisted between the ecologically dissimilar co-occurring species (BM–LTM) but were not detected between the ecologically similar co-occurring species (BM–RM), consistent with an influence of ecological similarity on microbial differentiation.

**Figure 5.**
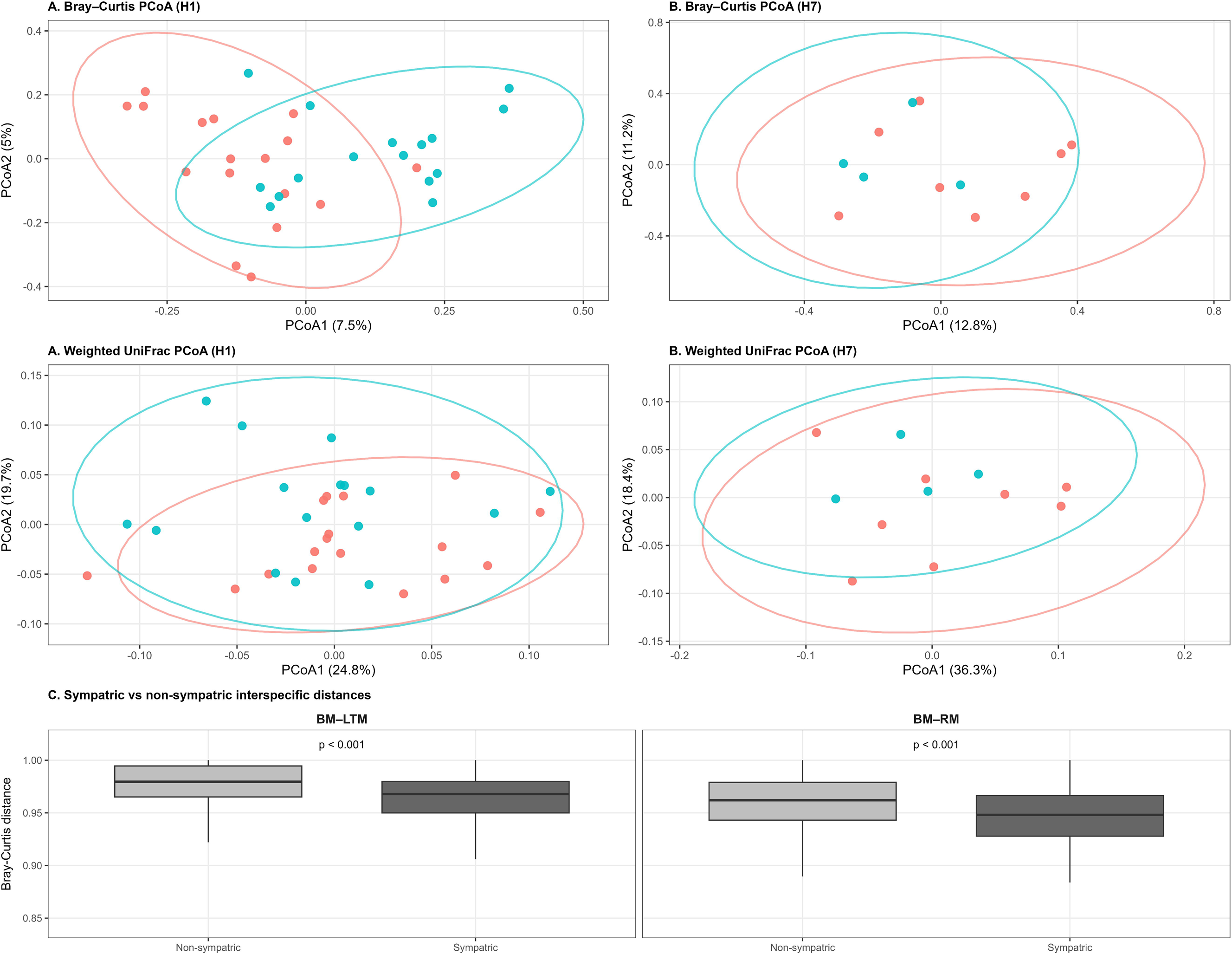
Microbiome similarity among sympatric macaque populations. Principal coordinates analysis (PCoA) of gut microbial communities in sympatric macaque populations occupying shared anthropogenic environments. **(A)** Bray–Curtis and weighted UniFrac ordinations comparing BM and LTM at site H1. **(B)** Bray–Curtis and weighted UniFrac ordinations comparing BM and RM at site H7. Ellipses represent 95% confidence intervals around host centroids. **(C)** Pairwise Bray–Curtis dissimilarities comparing sympatric and non-sympatric BM– LTM and BM–RM species pairs.

To further evaluate the influence of sympatry, we compared pairwise interspecific Bray–Curtis distances between sympatric and non-sympatric species pairs. In both BM–LTM and BM–RM comparisons, sympatric pairs exhibited modestly lower microbiome dissimilarities than non-sympatric pairs (BM–LTM: 0.963 vs. 0.977; BM–RM: 0.947 vs. 0.959), indicating a small increase in gut microbial similarity associated with site co-occurrence rather than a substantial shift in composition. However, substantial interspecific differentiation persisted even among sympatric hosts, and BM–RM pairs remained consistently more similar than BM–LTM pairs. To evaluate this pattern more broadly, pairwise Bray–Curtis distances across all samples were classified into three comparison categories: sympatric heterospecific pairs (different species, same site), non-sympatric heterospecific pairs (different species, different sites), and conspecific pairs from different sites. A Kruskal–Wallis test indicated that dissimilarity distributions differed significantly among these categories (χ² = 266.99, df = 2, p < 2.2×10 ¹). Pairwise Wilcoxon tests (BH-corrected) showed that sympatric heterospecific pairs and conspecific pairs from different sites did not differ significantly from one another (p = 0.72), while non-sympatric heterospecific pairs showed significantly greater dissimilarity than both sympatric heterospecific pairs (p = 1.8×10 ¹) and conspecific pairs from different sites (p < 2×10 ¹). Consistent with the pairwise comparisons above, sympatric heterospecific pairs showed Bray–Curtis dissimilarities statistically indistinguishable from conspecific pairs sampled at different sites, and significantly lower than non-sympatric heterospecific pairs (Fig. 5C). Co-occurrence was therefore associated with a measurable, though modest, increase in interspecific microbiome similarity, bringing sympatric heterospecific pairs into the range of variation observed among conspecifics sampled at different sites.

Despite this compositional divergence across species and ecological contexts, functional profiling of the microbiome revealed a different pattern: broad conservation of pathway architecture rather than differentiation.

### Microbial functional composition across habitat categories

Shotgun metagenomic sequencing to characterize functional pathway profiles showed substantial overlap in functional composition across macaque hosts and habitat categories (Fig. 6). Principal coordinates analysis of Bray–Curtis distances showed substantial overlap among host species with PCoA1 explaining 51.6% of the total variation. PERMANOVA analyses detected no significant functional differentiation across host species and habitat categories (R² = 0.347, F = 1.35, p = 0.126). Similarly, multivariate dispersion did not differ significantly between host species (p = 0.233) or anthropogenic environments (p = 0.402), indicating comparable levels of functional heterogeneity across groups (Fig. 6A).

**Figure 6.**
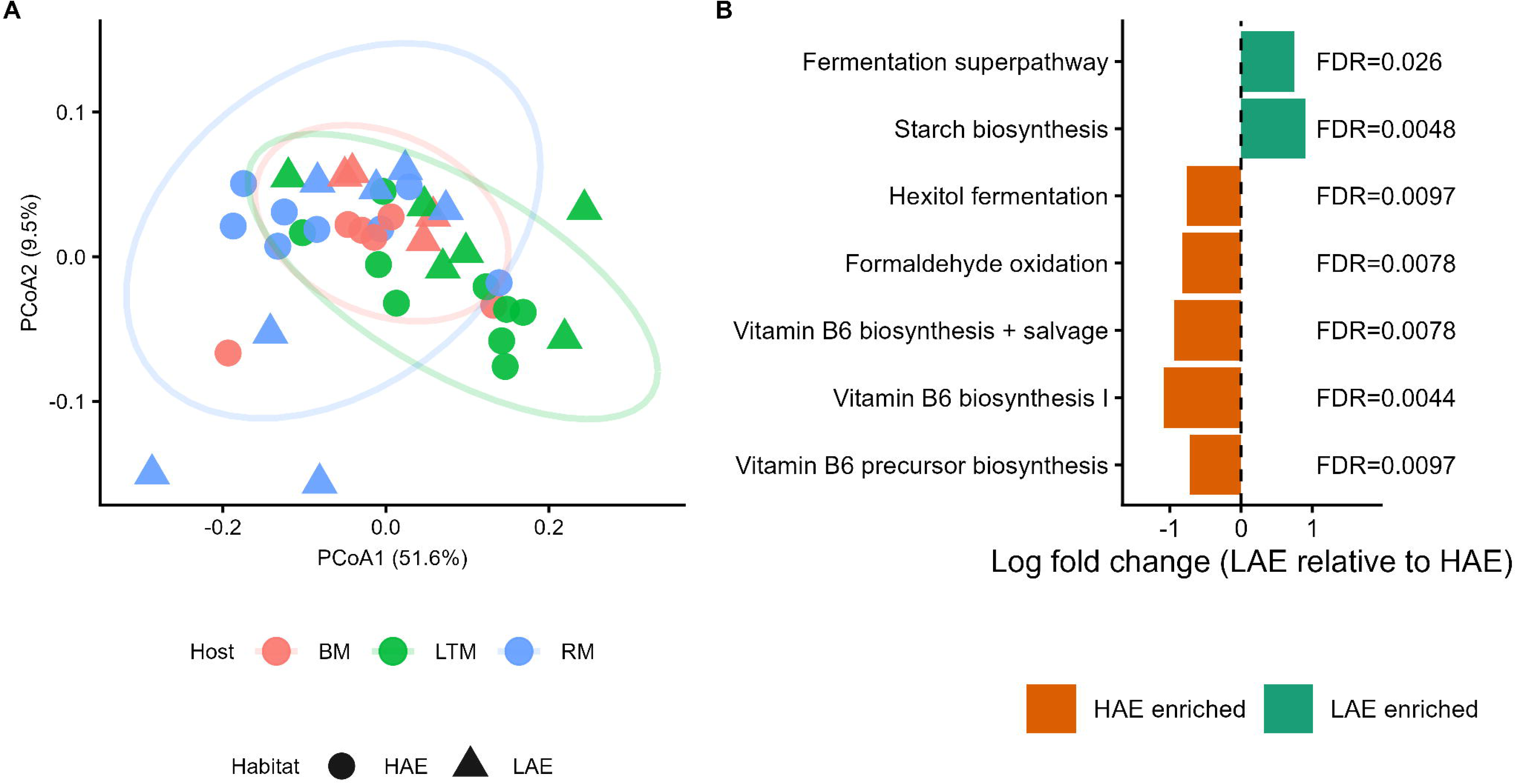
Functional structure and differential pathway enrichment across habitat categories. (A) PCoA of Bray–Curtis dissimilarities based on microbial functional pathway profiles. Colours indicate host species (BM; LTM; RM) and symbols indicate anthropogenic environments (LAE, low anthropogenic exposure; HAE, high anthropogenic exposure). (B) Differentially abundant pathways between LAE and HAE habitats identified using ANCOM-BC2. Log fold changes are expressed relative to HAE habitats as the reference level: positive values indicate higher relative abundance in LAE habitats, and negative values indicate higher relative abundance in HAE habitats. FDR-adjusted q-values from ANCOM-BC2 are shown for each pathway.

Differential functional pathway analyses identified selective restructuring of metabolic functions across anthropogenic exposure categories (Fig. 6B). High-anthropogenic-exposure habitats were enriched in pathways associated with pyridoxal 5-phosphate (vitamin B6) biosynthesis and salvage, formaldehyde oxidation, and hexitol fermentation. In contrast, low-anthropogenic-exposure habitats were enriched in pathways related to starch biosynthesis and a fermentation superpathway. These results indicate that, despite substantial overlap in overall functional composition, specific metabolic pathways exhibited exposure-associated shifts in relative abundance.

## Discussion

### Host ecology as a filter on microbiome responses to anthropogenic change

The magnitude and nature of microbiome restructuring differed among the three macaque species. BM and RM, both classified as synurbanized, showed comparatively modest restructuring, whereas LTM, a forest specialist, showed more extensive restructuring. This pattern is consistent with the general expectation that habitat specialists are more sensitive to environmental disturbance than ecological generalists (Clavel, Julliard and Devictor, 2011; Barelli *et al*., 2015; Bista *et al*., 2021). The three species also differed in which biological dimension of the microbiome carried the response: BM exhibited restructuring primarily through compositional turnover without substantial diversity loss, RM responses were concentrated in shifts among specific taxa rather than broad diversity change, and LTM showed both diversity loss and phylogenetically structured compositional change.

These species-specific response modes indicate that host ecology shapes not only whether microbiomes respond to anthropogenic environments, but which biological dimension carries that response. Ecologically flexible species may buffer microbiome disruption in response to anthropogenic pressure, and species sharing the same landscape may respond differently to identical anthropogenic exposure depending on their degree of ecological flexibility (Barelli *et al*., 2020; Fackelmann *et al*., 2021; Wasimuddin *et al*., 2022; Muhammad *et al*., 2023). Experimental work further supports this pattern under controlled conditions: a dietary specialist exposed to an identical sequence of environmental stressors showed substantially more variable and directional microbiome responses than a co-exposed generalist, indicating that ecological specialization predicts microbiome sensitivity even independent of field-based confounds (Koziol *et al*., 2023). Our results extend this pattern beyond within-landscape comparisons to species differing more fundamentally in ecological history, suggesting that ecological flexibility governs not just whether microbiome disruption occurs but which biological dimension it manifests (Ley *et al*., 2008; Moran, Ochman and Hammer, 2019; Greene *et al*., 2020; Mallott and Amato, 2021; Araujo *et al*., 2025) .

### The lion-tailed macaque as a sentinel system

LTM exhibited the strongest microbiome response to anthropogenic exposure, characterized by reduced diversity and extensive phylogenetically structured compositional change. These patterns are broadly consistent with expectations for a habitat specialist encountering environmental conditions outside its typical ecological range (Devictor, Julliard and Jiguet, 2008; Clavel, Julliard and Devictor, 2011; Bista *et al*., 2021). Unlike the synurbanized species, whose microbiomes exhibited comparatively modest restructuring, LTM showed evidence of broader taxonomic restructuring that may reflect a greater sensitivity of specialist-associated microbiomes to anthropogenic environmental change.

From a conservation perspective, this distinction may be important. LTM is an endangered rainforest specialist with a restricted distribution and high sensitivity to habitat disturbance (Dhawale, Kumar and Sinha, 2020; Mahato *et al*., 2026). These microbiome shifts may reflect altered fiber-processing capacity and other functional changes associated with dietary and environmental change (Barelli *et al*., 2015; Moustafa *et al*., 2021; Wasimuddin *et al*., 2022), capturing a physiological response to anthropogenic disturbance. We therefore suggest that microbiome profiling, particularly of microbial groups associated with plant fiber degradation, may have potential as an additional indicator of ecological disturbance in fragmented LTM populations, capable of detecting physiological stress before it manifests as measurable changes in behaviour, body condition, or the population-level demographic shifts already used in conservation monitoring (Mahato *et al*., 2026), consistent with the broader potential for microbiome-based conservation indicators (Trevelline *et al*., 2019). Whether these microbiome changes are reversible remains an important question for future conservation research.

### Functional redundancy as a mechanism of compositional–functional decoupling

One of the most notable findings of this study was the apparent decoupling between taxonomic restructuring and functional composition. Despite substantial taxonomic and phylogenetic change across species and anthropogenic exposure categories, functional profiles remained comparatively conserved. This pattern is consistent with functional redundancy in microbial communities (Louca *et al*., 2018; Rennison, Rudman and Schluter, 2019), whereby alternative microbial taxa with overlapping metabolic capabilities maintain similar community-level functions despite changes in taxonomic membership. Taxa enriched and depleted under anthropogenic exposure largely occupy overlapping metabolic roles, particularly in carbohydrate fermentation, short-chain fatty acid production, and vitamin biosynthesis. This suggests that the functional roles associated with declining taxa may be compensated for by taxa that increase in abundance, thereby preserving broad metabolic capabilities despite substantial shifts in community composition (Adair and Douglas, 2017; Li *et al*., 2024). The absence of shared differentially abundant ASVs across host species, combined with conservation of functional pathways, further suggests that functionally equivalent outcomes can be achieved through taxonomically distinct routes. In this framework, host ecological differences may shape the taxonomic pathways leading to broadly conserved functional potential.

The selective enrichment of pathways related to vitamin B6 biosynthesis, formaldehyde oxidation, and hexitol fermentation under HAE conditions deserves specific consideration because it may indicate the limits of this redundancy. Vitamin B6 enrichment could reflect microbial compensation for altered dietary nutrient availability under anthropogenic feeding regimes. Alternatively, it may result from selection for microbial taxa adapted to simple-carbohydrate-rich diets (Mayengbam, Chleilat and Reimer, 2020; Wan *et al*., 2022; Wibowo and Pramadhani, 2024; Tarracchini *et al*., 2025). Similarly, enrichment of hexitol fermentation pathways may reflect shifts in dietary carbohydrate availability, which is known to influence gut microbial composition and metabolic activity (Zong *et al*., 2024). Conversely, pathways associated with starch fermentation under LAE conditions are consistent with greater reliance on complex plant-derived carbohydrates in forest-associated populations (Warren *et al*., 2018). These patterns suggest that while overall functional architecture remains conserved, the metabolic emphasis within that architecture may shift according to ecological context. Resolving whether these differences represent changes in functional potential or actual metabolic activity will require metatranscriptomic approaches.

### Sympatry and ecological context

The sympatric analyses provide important insight into the relative contributions of ecological context and host identity. Sympatric co-occurrence increased microbiome similarity, indicating that local ecological conditions can partially offset host-associated differentiation. This finding engages directly with the long-standing debate between phylosymbiosis and ecological lability in host-associated microbiomes (Mazel *et al*., 2018; Moeller and Sanders, 2020). The present data support neither framework exclusively. Although sympatric co-occurrence increased microbiome similarity, host-associated differences persisted, and the species pairs that remained most similar after accounting for shared habitat were those that were also most similar in their ecological integration into anthropogenic environments — sharing a site itself did not predict residual similarity as strongly as shared ecology did (Fu *et al*., 2021; Liu *et al*., 2021; Anders *et al*., 2022; Heni *et al*., 2023; Tuoliu *et al*., 2024). Species that were more similar in their ecological integration into anthropogenic environments exhibited greater microbiome similarity than ecologically divergent species occupying the same sites.

These findings suggest that ecological context and host ecology should not be treated as competing explanations for microbiome structure. Instead, ecological context determines the environmental signals available to microbial communities, while host ecological and evolutionary characteristics determine how strongly those signals are expressed within the microbiome. Sympatric co-occurrence reduces variation attributable to environmental context, thereby revealing the residual contribution of host ecology more clearly. At the two sympatric sites, species-associated microbiome differences persisted between the ecologically dissimilar co-occurring species (BM and LTM) but were undetectable between the ecologically similar co-occurring species (BM and RM), indicating that co-occurrence reduces microbial differentiation only when the co-occurring hosts are themselves ecologically similar. Contrasting outcomes at the two sympatric sites further suggest that microbiome similarity depends on ecological similarity rather than co-occurrence itself.

### Implications for understanding wildlife microbiome responses to environmental change

Taken together, the findings of this study support a framework in which anthropogenic environmental change does not generate a common microbiome response across wildlife, but rather interacts with host ecology to produce a spectrum of outcomes whose nature and magnitude reflect the ecological and evolutionary history of each host (Ley *et al*., 2008; Alberdi *et al*., 2016; Flynn *et al*., 2022; Kuthyar *et al*., 2022; Wasimuddin *et al*., 2022; Wills *et al*., 2022; Araujo *et al*., 2025). This framing has several implications for how microbiome studies of wildlife responses to environmental change should be designed and interpreted.

First, single-species studies may not reliably distinguish host-specific from environmentally general microbiome responses (Moy, Diakiw and Amato, 2023; Amato *et al*., 2025); comparative frameworks spanning species that differ in ecological flexibility but share anthropogenic exposure gradients are needed to partition these effects. Second, our results show that restricting analysis to a single microbiome dimension risks missing the complete pattern of restructuring. Alpha diversity in isolation would have identified only LTM as responding to anthropogenic exposure. Beta diversity analyses, however, revealed significant habitat-associated differentiation across all three species. Differential abundance and functional analyses then added resolution unavailable from either metric alone. Third, functional composition should not be assumed to covary with taxonomic composition (Moya and Ferrer, 2016; Inkpen *et al*., 2017; Louca *et al*., 2018): the decoupling observed here, and the identification of specific pathway-level shifts within a conserved functional architecture, illustrates that taxonomic restructuring and functional change are partly independent axes of microbiome response to environmental pressure (Moya and Ferrer, 2016; Sambamoorthy and Raman, 2018; Cross *et al*., 2025; John *et al*., 2026). The finding that functional gene composition is conserved across anthropogenic gradients while specific pathways shift in relative abundance raises an important distinction between functional potential and functional activity that current shotgun metagenomic approaches cannot fully resolve. Differential expression of shared metabolic pathways, independent of gene presence itself, can meaningfully shape microbiome function (Ho and Huang, 2025). This distinction may be particularly relevant to how microbiome-mediated physiology adjusts to anthropogenic environments, a possibility our gene-level data cannot resolve. Metatranscriptomic analyses across anthropogenic gradients in systems such as this one represent a natural next step for resolving whether the functional buffering apparent at the gene composition level translates into genuinely similar metabolic output. Fourth, microbiome responses to anthropogenic change are inherently scale-dependent, and analyses restricted to a single level of biological organization will systematically underestimate the entirety of restructuring signals (Ladau and Eloe-Fadrosh, 2019; Stothart and Newman, 2021; Medeiros *et al*., 2026). The present study highlights this across three scales simultaneously: (i) individual-level dispersion heterogeneity revealed within-species variability in anthropogenic response that community-level analyses would otherwise have obscured, (ii) species-level comparisons revealed divergent restructuring modes, from diversity-stable compositional reorganization in BM to community-wide diversity loss in LTM, that single-species designs cannot detect, and (iii) interspecific sympatric analyses revealed that the relative contributions of ecological context and host identity are only detectable when within-site and across-site comparisons are examined together. Explicitly incorporating scale as an analytical dimension while examining microbiome responses at individual, species, and interspecific levels simultaneously reveals restructuring signals that single-scale analyses may miss.

### Limitations and future directions

As with most field-based wildlife microbiome studies, our findings should be interpreted within the context of several practical constraints. First, anthropogenic exposure was classified into two broad categories to allow consistent comparisons across species and sites. Because human influence occurs along a continuum, some site effects are inevitably linked with host differences, particularly in comparisons among species. Future studies incorporating continuous measures of anthropogenic exposure could help separate these effects more clearly. Second, this study provides a cross-sectional view of the gut microbiome and cannot determine whether the reduced microbial diversity observed in lion-tailed macaques represents a temporary or persistent change. Longitudinal sampling would help address this question. Although a small number of rhesus macaque samples may have come from the same individual, such cases are expected to be rare and are unlikely to affect the overall conclusions. Finally, our functional analyses describe the genetic potential of the microbiome rather than its activity. Metatranscriptomic approaches would be needed to determine whether the conserved pathways identified here are expressed similarly across hosts and habitats.

## Conclusions

The gut microbiome does not respond consistently to anthropogenic change across host species. Instead, microbiome responses are shaped by each species’ ecological and evolutionary history. The three macaque species examined here illustrate this pattern, with responses ranging from compositional reorganization while maintaining microbial diversity in a long-synurbanized generalist to reduced diversity and pronounced phylogenetic restructuring in a forest specialist. Despite these divergent taxonomic responses, the functional potential of the gut microbiome remained largely conserved, suggesting that microbial communities can maintain similar metabolic capabilities even when their taxonomic composition is altered. Together, these findings highlight the importance of comprehensively characterizing both the taxonomic and functional dimensions of the gut microbiome to understand the consequences of anthropogenic change for host adaptation and health, which remain important questions in wildlife ecology, host–microbiome biology, and conservation science.

## Acknowledgements

We sincerely thank the Forest Departments of Karnataka, Telangana, and Tamil Nadu for granting the necessary research permits and for their continued support throughout this study. We are particularly grateful to the forest officers and frontline staff in all three states for their invaluable assistance in locating macaque troops, facilitating access to field sites, supporting sample collection, and providing logistical support, including local accommodation and field coordination at several study sites. We thank Vignesh for assistance with field sampling in Valparai, Ashish Eradath in Mannanur, Prabhu in Chincholi, and ShamSundar Rao in Agumbe. Their support was invaluable to the successful completion of the fieldwork. We also thank the members of the Praveen Karanth Laboratory for their assistance with routine laboratory work during sample processing.

## Data Accessibility Statement

Raw 16S rRNA gene amplicon and shotgun metagenomic sequence data generated in this study are deposited with the Indian Biological Data Centre (IBDC) under Study accession PRJINA00733 and with the Indian Nucleotide Data Archive (INDA) under accession INRP000715. The data and associated sample metadata will be made publicly available upon publication.

## Author Contributions

V.K. conceived and designed the study, conducted fieldwork and laboratory research, performed the bioinformatic and statistical analyses, interpreted the data, and wrote the manuscript. S.R. contributed to study design, interpretation of the results, and critically revised the manuscript. P.K. contributed to interpretation of the results, provided intellectual input during manuscript preparation, and critically revised the manuscript.

## Ethics Statement

Fieldwork was approved by the Institutional Ethics Committee of **the** National Institute of Advanced Studies (NIAS) and was conducted in accordance with applicable institutional ethical guidelines and permissions from the relevant state authorities. Fecal samples were collected non-invasively from free-ranging macaques without capturing, handling, restraining, or otherwise disturbing the animals. Field sampling was conducted under permissions issued by the respective State Forest Departments of Karnataka (No. KFD/WL/E2(RE)25/2024), Telangana (Rc.No.11853/2017/WL-2 (ii); File No. PCCF-WL2/WL19/3/2022-WILD LIFE SECTION), and Tamil Nadu (Proceeding No. WL5(A)/9635/2024; Permission No. 30/2024).

## References

Adair, K.L. and Douglas, A.E. (2017) “Making a microbiome: the many determinants of host-associated microbial community composition,” Current Opinion in Microbiology, 35, pp. 23–29. Available at: 10.1016/j.mib.2016.11.002.

Adair, M.G. et al. (2025) “Anthropogenic reverberations on the gut microbiome of dwarf chameleons (Bradypodion),” PeerJ, 13, p. e18811. Available at: 10.7717/peerj.18811.

Alberdi, A. et al. (2016) “Do Vertebrate Gut Metagenomes Confer Rapid Ecological Adaptation?,” Trends in Ecology & Evolution, 31(9), pp. 689–699. Available at: 10.1016/j.tree.2016.06.008.

Amato, K.R. et al. (2016) “Phylogenetic and ecological factors impact the gut microbiota of two Neotropical primate species,” Oecologia, 180(3), pp. 717–733. Available at: 10.1007/s00442-015-3507-z.

Amato, K.R. et al. (2025) “Supplementation With Human Foods Affects the Gut Microbiota of Wild Howler Monkeys,” American Journal of Primatology, 87(4), p. 70029.

Anders, J.L. et al. (2022) “Dietary niche breadth influences the effects of urbanization on the gut microbiota of sympatric rodents,” Ecology and Evolution, 12(9), p. e9216. Available at: 10.1002/ece3.9216.

Araujo, G. et al. (2025) “A mechanistic framework for complex microbe-host symbioses,” Trends in Microbiology, 33(1), pp. 96–111. Available at: 10.1016/j.tim.2024.08.002.

Barelli, C. et al. (2015) “Habitat fragmentation is associated to gut microbiota diversity of an endangered primate: implications for conservation,” Scientific Reports, 5(1), p. 14862. Available at: 10.1038/srep14862.

Barelli, C., et al. (2020) “The gut microbiota communities of wild arboreal and ground-feeding tropical primates are affected differently by habitat disturbance,” *Msystems*, 5(3), pp. 10–1128.

Barko, P.C. et al. (2018) “The Gastrointestinal Microbiome: A Review,” Journal of Veterinary Internal Medicine, 32(1), pp. 9–25. Available at: 10.1111/jvim.14875.

Beeby, N., Pierre, L.J. and Guy, R.F.J. (2025) “Climate, diet, and nutrition drive gut microbiome variation in a fruit-specialist primate,” Sci Rep, 15, p. 21110. Available at: 10.1038/s41598-025-07399-3.

Bista, D. et al. (2021) “Movement and dispersal of a habitat specialist in human-dominated landscapes: a case study of the red panda,” Movement Ecology, 9, p. 62. Available at: 10.1186/s40462-021-00297-z.

Bolyen, E. et al. (2019) “Reproducible, interactive, scalable and extensible microbiome data science using QIIME 2,” Nature Biotechnology, 37(8), pp. 852–857. Available at: 10.1038/s41587-019-0209-9.

Callahan, B.J. et al. (2016) “DADA2: High-resolution sample inference from Illumina amplicon data,” Nature Methods, 13(7), pp. 581–583. Available at: 10.1038/nmeth.3869.

Chen, T. et al. (2020) “Gut microbiota of provisioned and wild rhesus macaques (Macaca mulatta) living in a limestone forest in southwest Guangxi, China,” MicrobiologyOpen, 9(3), p. e981. Available at: 10.1002/mbo3.981.

Clavel, J., Julliard, R. and Devictor, V. (2011) “Worldwide decline of specialist species: toward a global functional homogenization?,” Frontiers in Ecology and the Environment, 9(4), pp. 222–228. Available at: 10.1890/080216.

Clayton, J.B. et al. (2016) “Captivity humanizes the primate microbiome,” Proceedings of the National Academy of Sciences, 113(37), pp. 10376–10381. Available at: 10.1073/pnas.1521835113.

Clayton, J.B. et al. (2018) “The gut microbiome of nonhuman primates: lessons in ecology and evolution,” American Journal of Primatology, 80(6), p. 22867.

Cross, K. et al. (2025) “Microbiome metabolic capacity is buffered against phylotype losses by functional redundancy,” Applied and Environmental Microbiology, 91(2), pp. e02368–24. Available at: 10.1128/aem.02368-24.

Degregori, S. et al. (2025) “Comparative gut microbiome research through the lens of ecology: theoretical considerations and best practices,” Biological Reviews, 100(2), pp. 748–763. Available at: 10.1111/brv.13161.

Devictor, V., Julliard, R. and Jiguet, F. (2008) “Distribution of specialist and generalist species along spatial gradients of habitat disturbance and fragmentation,” Oikos, 117(4), pp. 507–514. Available at: 10.1111/j.0030-1299.2008.16215.x.

Dhawale, A.K., Kumar, M.A. and Sinha, A. (2020) “Changing ecologies, shifting behaviours: Behavioural responses of a rainforest primate, the lion-tailed macaque Macaca silenus, to a matrix of anthropogenic habitats in southern India,” PLoS One, 15(9), p. e0238695.

Di Rienzi, S.C. and Britton, R.A. (2020) “Adaptation of the Gut Microbiota to Modern Dietary Sugars and Sweeteners,” Advances in Nutrition, 11(3), pp. 616–629. Available at: 10.1093/advances/nmz118.

Dillard, B.A., et al. (2022) “Humanization of wildlife gut microbiota in urban environments,” eLife. Edited by P.J. Turnbaugh et al., 11, p. e76381. Available at: 10.7554/eLife.76381.

Erinjery, J.J. et al. (2017) “Losing its ground: A case study of fast declining populations of a ‘least-concern’ species, the bonnet macaque (Macaca radiata),” PLOS ONE, 12(8), p. e0182140. Available at: 10.1371/journal.pone.0182140.

Fackelmann, G. et al. (2021) “Human encroachment into wildlife gut microbiomes,” Communications Biology, 4(1), p. 800. Available at: 10.1038/s42003-021-02315-7.

Flynn, J.K. et al. (2022) “Host Genetics and Environment Shape the Composition of the Gastrointestinal Microbiome in Nonhuman Primates,” Microbiology Spectrum, 11(1), pp. e02139–22. Available at: 10.1128/spectrum.02139-22.

Fu, H. et al. (2021) “Sympatric Yaks and Plateau Pikas Promote Microbial Diversity and Similarity by the Mutual Utilization of Gut Microbiota,” Microorganisms, 9(9). Available at: 10.3390/microorganisms9091890.

Gloor, G. (2023) *ANOVA-Like Differential Expression tool for high throughput sequencing data - ALDEx2*, *R-universe*. Available at: https://bioc.r-universe.dev/articles/ALDEx2/ALDEx2_vignette.html (Accessed: June 25, 2026).

Greene, L.K. et al. (2020) “A role for gut microbiota in host niche differentiation,” The ISME Journal, 14(7), pp. 1675–1687. Available at: 10.1038/s41396-020-0640-4.

Heni, A.C. et al. (2023) “Wildlife gut microbiomes of sympatric generalist species respond differently to anthropogenic landscape disturbances,” Animal Microbiome, 5(1), p. 22. Available at: 10.1186/s42523-023-00237-9.

Ho, P.-Y. and Huang, K.C. (2025) “Challenges in interpreting functional redundancy and quantifying functional selection in microbial communities,” Cell Systems, 16(8), p. 101350. Available at: 10.1016/j.cels.2025.101350.

Inkpen, S.A. et al. (2017) “The coupling of taxonomy and function in microbiomes,” Biology & Philosophy, 32(6), pp. 1225–1243. Available at: 10.1007/s10539-017-9602-2.

John, J. et al. (2026) “Functional redundancy and metabolic flexibility of microbial communities in two Mid-Atlantic bays,” ISME Communications, 6(1), p. ycag021. Available at: 10.1093/ismeco/ycag021.

Koziol, A. et al. (2023) “Mammals show distinct functional gut microbiome dynamics to identical series of environmental stressors,” mBio, 14(5), pp. e01606–23. Available at: 10.1128/mbio.01606-23.

Kumar, R., Radhakrishna, S. and Sinha, A. (2011) “Of Least Concern? Range Extension by Rhesus Macaques (Macaca mulatta) Threatens Long-Term Survival of Bonnet Macaques (M. radiata) in Peninsular India,” International Journal of Primatology, 32(4), pp. 945–959. Available at: 10.1007/s10764-011-9514-y.

Kuthyar, S. et al. (2022) “Limited microbiome differences in captive and semi-wild primate populations consuming similar diets,” FEMS Microbiology Ecology, 98(10), p. fiac098. Available at: 10.1093/femsec/fiac098.

Ladau, J. and Eloe-Fadrosh, E.A. (2019) “Spatial, Temporal, and Phylogenetic Scales of Microbial Ecology,” Trends in Microbiology, 27(8), pp. 662–669. Available at: 10.1016/j.tim.2019.03.003.

Lee, W. et al. (2019) “Gut microbiota composition of Japanese macaques associates with extent of human encroachment,” American Journal of Primatology, 81(12), p. e23072. Available at: 10.1002/ajp.23072.

Ley, R.E. et al. (2008) “Evolution of Mammals and Their Gut Microbes,” Science, 320(5883), pp. 1647–1651. Available at: 10.1126/science.1155725.

Li, Y. et al. (2024) “The role of gut microbiota in a generalist, golden snub-nosed monkey, adaptation to geographical diet change,” Animal Microbiome, 6(1), p. 63. Available at: 10.1186/s42523-024-00349-w.

Lin, H. and Peddada, S.D. (2024) “Multigroup analysis of compositions of microbiomes with covariate adjustments and repeated measures,” Nature Methods, 21(1), pp. 83–91. Available at: 10.1038/s41592-023-02092-7.

Littleford-Colquhoun, B.L. et al. (2019) “City life alters the gut microbiome and stable isotope profiling of the eastern water dragon (Intellagama lesueurii).” Available at: 10.1111/mec.15240.

Liu, R. et al. (2021) “Fecal Bacterial Community of Allopatric Przewalski’s Gazelles and Their Sympatric Relatives,” Frontiers in Microbiology, 12. Available at: 10.3389/fmicb.2021.737042.

Liu, X. et al. (2024) “Comparing the gut microbiota of Sichuan golden monkeys across multiple captive and wild settings: roles of anthropogenic activities and host factors,” BMC Genomics, 25(1), p. 148.

Louca, S. et al. (2018) “Function and functional redundancy in microbial systems,” Nature Ecology & Evolution, 2(6), pp. 936–943. Available at: 10.1038/s41559-018-0519-1.

Ma, Z. (Sam) (2026) “How Host Phylogeny and Diet Shape the Specificity and Specificity Diversity of Animal Gut Microbiomes,” Environmental Microbiology Reports, 18(1), p. e70253. Available at: 10.1111/1758-2229.70253.

Mahato, S. et al. (2026) “Differential demographic responses of lion-tailed macaques to habitat fragmentation: Four decades of population monitoring in the Anamalai Hills, Western Ghats and perspectives for management and conservation,” Journal for Nature Conservation, 91, p. 127236. Available at: 10.1016/j.jnc.2026.127236.

Mallott, E.K. and Amato, K.R. (2021) “Host specificity of the gut microbiome,” Nature Reviews Microbiology, 19(10), pp. 639–653. Available at: 10.1038/s41579-021-00562-3.

Maraci, Ö. et al. (2022) “Changes to the gut microbiota of a wild juvenile passerine in a multidimensional urban mosaic,” Scientific Reports, 12(1), p. 6872. Available at: 10.1038/s41598-022-10734-7.

Marsh, K.J., Bearhop, S. and Harrison, X.A. (2024) “Linking microbiome temporal dynamics to host ecology in the wild,” Trends in Microbiology, 32(11), pp. 1060–1071. Available at: 10.1016/j.tim.2024.05.001.

Mayengbam, S., Chleilat, F. and Reimer, R.A. (2020) “Dietary Vitamin B6 Deficiency Impairs Gut Microbiota and Host and Microbial Metabolites in Rats,” Biomedicines, 8(11). Available at: 10.3390/biomedicines8110469.

Mazel, F. et al. (2018) “Is Host Filtering the Main Driver of Phylosymbiosis across the Tree of Life?,” mSystems, 3(5), p. 10.1128/msystems.00097-18. Available at: 10.1128/msystems.00097-18.

McDonald, J.E., Marchesi, J.R. and Koskella, B. (2020) “Application of ecological and evolutionary theory to microbiome community dynamics across systems,” Proceedings of the Royal Society B: Biological Sciences, 287(1941), p. 20202886. Available at: 10.1098/rspb.2020.2886.

McMurdie, P.J. and Holmes, S. (2013) “phyloseq: an R package for reproducible interactive analysis and graphics of microbiome census data,” PloS One, 8(4), p. e61217. Available at: 10.1371/journal.pone.0061217.

Medeiros, M.J. et al. (2026) “Microbiome Structure of Endemic Hawaiian Drosophila Is Shaped More by Habitat Than Host Identity,” Molecular Ecology, 35(11), p. e70417. Available at: 10.1111/mec.70417.

Moeller, A.H. et al. (2013) “Sympatric chimpanzees and gorillas harbor convergent gut microbial communities,” Genome Research, 23(10), pp. 1715–1720. Available at: 10.1101/gr.154773.113.

Moeller, A.H. and Sanders, J.G. (2020) “Roles of the gut microbiota in the adaptive evolution of mammalian species,” Philosophical Transactions of the Royal Society B: Biological Sciences, 375(1808), p. 20190597. Available at: 10.1098/rstb.2019.0597.

Moran, N.A., Ochman, H. and Hammer, T.J. (2019) “Evolutionary and Ecological Consequences of Gut Microbial Communities,” Annual Review of Ecology, Evolution, and Systematics, 50(Volume 50, 2019), pp. 451–475. Available at: 10.1146/annurev-ecolsys-110617-062453.

Moustafa, M.A.M. et al. (2021) “Anthropogenic interferences lead to gut microbiome dysbiosis in Asian elephants and may alter adaptation processes to surrounding environments,” Scientific Reports, 11(1), p. 741. Available at: 10.1038/s41598-020-80537-1.

Moy, M., Diakiw, L. and Amato, K.R. (2023) “Human-influenced diets affect the gut microbiome of wild baboons,” Scientific Reports, 13(1), p. 11886. Available at: 10.1038/s41598-023-38895-z.

Moya, A. and Ferrer, M. (2016) “Functional Redundancy-Induced Stability of Gut Microbiota Subjected to Disturbance,” Trends in Microbiology, 24(5), pp. 402–413. Available at: 10.1016/j.tim.2016.02.002.

Muhammad, R. et al. (2023) “Comparative analysis of gut microbiota between common (Macaca fascicularis fascicularis) and Burmese (M. f. aurea) long-tailed macaques in different habitats,” Scientific Reports, 13(1), p. 14950. Available at: 10.1038/s41598-023-42220-z.

Naik, T. et al. (2023) “High-quality single amplicon sequencing method for illumina MiSeq platform using pool of ‘N’ (0–10) spacer-linked target specific primers without PhiX spike-in,” BMC Genomics, 24(1), p. 141. Available at: 10.1186/s12864-023-09233-4.

Nguyen, H.K. et al. (2024) “Wildlife microbiomes and the city: a systematic review of urban impacts on wildlife bacterial communities,” Microbiota and Host, 2(1). Available at: 10.1530/MAH-24-0003.

Nguyen, J., Lara-Gutiérrez, J. and Stocker, R. (2021) “Environmental fluctuations and their effects on microbial communities, populations and individuals,” FEMS Microbiology Reviews, 45(4), p. fuaa068. Available at: 10.1093/femsre/fuaa068.

Peng, Y. et al. (2025) “Double-Edged Sword: Urbanization and Response of Amniote Gut Microbiome in the Anthropocene,” Microorganisms, 13(8). Available at: 10.3390/microorganisms13081736.

R Core Team (2025) “R: A language and environment for statistical computing (Version 4.5.2) [Computer software]. R Foundation for Statistical Computing. https://www.R-project.org/.”

Radhakrishna, S., Anand, S. and Sengupta, A. (2025) “Differing Trajectories: Understanding Coexistence Through the Human–Macaque Interface in India,” International Journal of Primatology, 46(6), pp. 1276–1290. Available at: 10.1007/s10764-025-00485-3.

Rennison, D.J., Rudman, S.M. and Schluter, D. (2019) “Parallel changes in gut microbiome composition and function during colonization, local adaptation and ecological speciation,” Proceedings of the Royal Society B: Biological Sciences, 286(1916), p. 20191911. Available at: 10.1098/rspb.2019.1911.

Requena, T., Martínez-Cuesta, M.C. and Peláez, C. (2018) “Diet and microbiota linked in health and disease,” Food & Function, 9(2), pp. 688–704. Available at: 10.1039/C7FO01820G.

Sambamoorthy, G. and Raman, K. (2018) “Understanding the evolution of functional redundancy in metabolic networks,” Bioinformatics, 34(17), pp. i981–i987. Available at: 10.1093/bioinformatics/bty604.

Sengupta, A., Anand, S. and Radhakrishna, S. (2025) “Behavioural Flexibility in Rhesus Macaques (Macaca mulatta) in Human-modified Environments,” Int J Primatol, 46, pp. 932–954. Available at: 10.1007/s10764-025-00506-1.

Sengupta, A. and Radhakrishna, S. (2016) “Influence of Fruit Availability on Fruit Consumption in a Generalist Primate, the Rhesus Macaque Macaca mulatta,” International Journal of Primatology, 37(6), pp. 703–717. Available at: 10.1007/s10764-016-9933-x.

Singh, M. (2019) “Management of Forest-Dwelling and Urban Species: Case Studies of the Lion-Tailed Macaque (Macaca silenus) and the Bonnet Macaque (M. radiata),” International Journal of Primatology, 40(6), pp. 613–629. Available at: 10.1007/s10764-019-00122-w.

Somers, S.E. et al. (2023) “Individual variation in the avian gut microbiota: The influence of host state and environmental heterogeneity,” Molecular Ecology, 32(12), pp. 3322–3339. Available at: 10.1111/mec.16919.

Stothart, M.R. and Newman, A.E.M. (2021) “Shades of grey: host phenotype dependent effect of urbanization on the bacterial microbiome of a wild mammal,” Animal Microbiome, 3(1), p. 46. Available at: 10.1186/s42523-021-00105-4.

Sugden, S., St. Clair, C.C. and Stein, L.Y. (2021) “Individual and Site-Specific Variation in a Biogeographical Profile of the Coyote Gastrointestinal Microbiota,” Microbial Ecology, 81(1), pp. 240–252. Available at: 10.1007/s00248-020-01547-0.

Tarracchini, C., et al. (2025) “Vitamin biosynthesis in the gut: interplay between mammalian host and its resident microbiota,” Microbiology and Molecular Biology Reviews. Edited by F. Rey, 89(2), pp. e00184–23. Available at: 10.1128/mmbr.00184-23.

Trevelline, B.K. et al. (2019) “Conservation biology needs a microbial renaissance: a call for the consideration of host-associated microbiota in wildlife management practices,” Proceedings of the Royal Society B: Biological Sciences, 286(1895), p. 20182448. Available at: 10.1098/rspb.2018.2448.

Tsuchida, S. et al. (2023) “The fecal microbiomes analysis of Marabou storks (Leptoptilos crumenifer) reveals their acclimatization to the feeding environment in the Kampala urban areas, Uganda,” Journal of Veterinary Medical Science, 85(4), pp. 450–458. Available at: 10.1292/jvms.22-0580.

Tuoliu, D. et al. (2024) “Bacterial microbiome and their assembly processing in two sympatric desert rodents (Dipus sagitta and Meriones meridianus) from different geographic sources,” Current Zoology, 71(4), pp. 440–448. Available at: 10.1093/cz/zoae062.

Wan, Z. et al. (2022) “Intermediate role of gut microbiota in vitamin B nutrition and its influences on human health,” Frontiers in Nutrition, 9. Available at: 10.3389/fnut.2022.1031502.

Warren, F.J., et al. (2018) “Food Starch Structure Impacts Gut Microbiome Composition,” mSphere. Edited by G. Suen, 3(3), pp. e00086–18. Available at: 10.1128/mSphere.00086-18.

Wasimuddin et al. (2022) “Anthropogenic Disturbance Impacts Gut Microbiome Homeostasis in a Malagasy Primate,” Frontiers in Microbiology, 13. Available at: 10.3389/fmicb.2022.911275.

Wibowo, S. and Pramadhani, A. (2024) “Vitamin B, Role of Gut Microbiota and Gut Health,” Vitamin B and Vitamin E - Pleiotropic and Nutritional Benefits. IntechOpen. Available at: 10.5772/intechopen.109485.

Wills, M.O. et al. (2022) “Host Species and Captivity Distinguish the Microbiome Compositions of a Diverse Zoo-Resident Non-Human Primate Population,” Diversity, 14(9). Available at: 10.3390/d14090715.

Xia, W. et al. (2022) “Functional convergence of Yunnan snub-nosed monkey and bamboo-eating panda gut microbiomes revealing the driving by dietary flexibility on mammal gut microbiome,” Computational and Structural Biotechnology Journal, 20, pp. 685–699. Available at: 10.1016/j.csbj.2022.01.011.

Yildirim, S. et al. (2010) “Characterization of the Fecal Microbiome from Non-Human Wild Primates Reveals Species Specific Microbial Communities,” PLOS ONE, 5(11), p. e13963. Available at: 10.1371/journal.pone.0013963.

Zong, X. et al. (2024) “Nondigestible carbohydrates and gut microbiota: A dynamic duo in host defense,” Animal Nutriomics, 1, p. e7. Available at: 10.1017/anr.2024.8.

