## Supplementary Information for "Host ecological context influences taxonomic diversity and functional conservation of gut microbiome across anthropogenic habitats in macaques"

**Figure S1.**

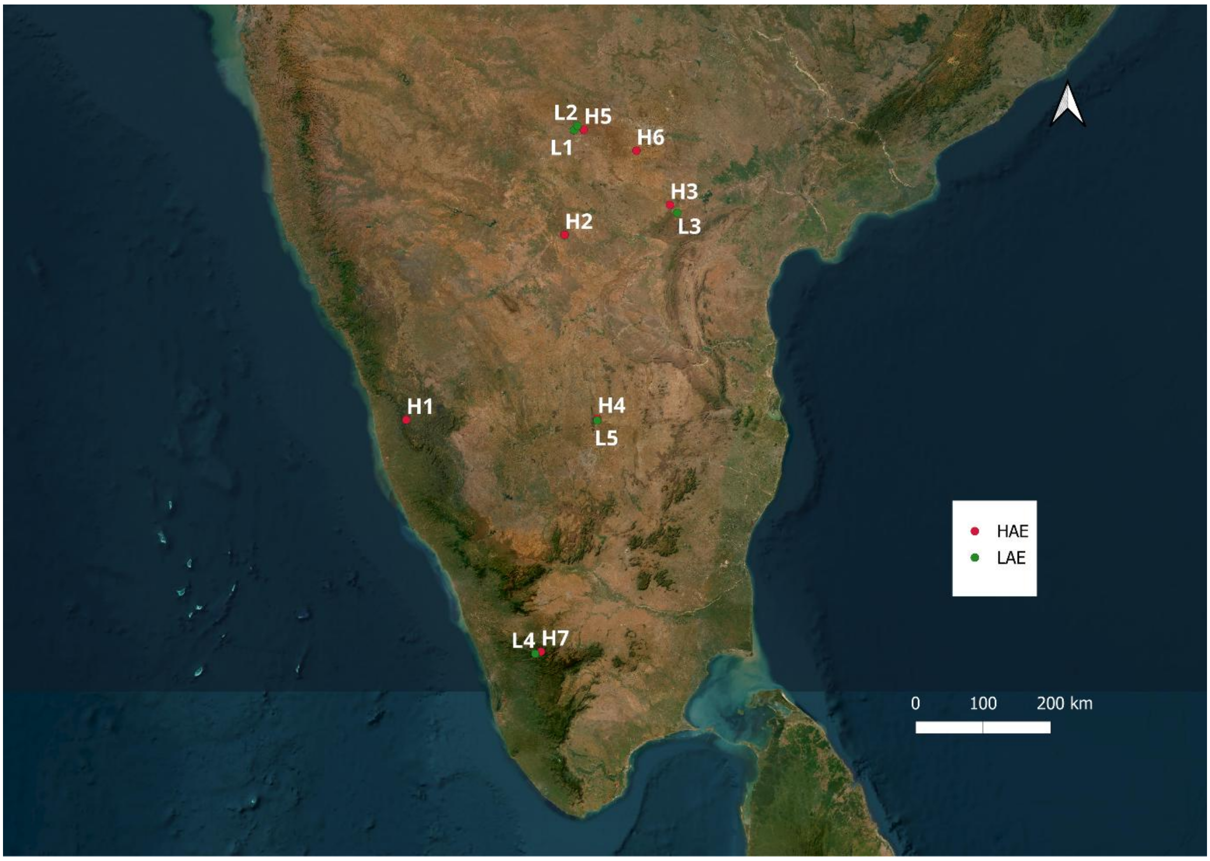


**Figure S1.** Geographic distribution of the 12 study sites across southern India included in this study. Sites are classified as **high anthropogenic exposure (HAE; red)** and **low anthropogenic exposure (LAE; green)** based on habitat characteristics, degree of human–macaque interaction, and access to anthropogenic food resources. Site codes (H1–H7 and L1–L5) correspond to the locations and site descriptions provided in **Table S1**.

**Table S1. Ecological characteristics of sampling sites and anthropogenic exposure classification.**

| Site | Host_species | N | Exposure | Habitat | Site_features | Food Access | Human_interaction | Sympatry |
|---|---|---|---|---|---|---|---|---|
| H1 | Bonnet macaque, Lion-tailed macaque | 33 | HAE | Forest | Roadside rainforest intersected by a major highway with continuous vehicular traffic and food discarded from vehicles | Provisioned | Low | Bonnet + Lion-tailed |
| H2 | Bonnet macaque | 8 | HAE | Rural | Rural settlement and agricultural landscape with regular kitchen raiding, crop access and provisioning | Provisioned | High | No |
| H3 | Bonnet macaque, Rhesus macaque | 19 | HAE | Rural | Village at the entrance of a wildlife sanctuary with tourist facilities, eateries and food waste providing regular anthropogenic food access | Provisioned | High | No |
| H4 | Bonnet macaque | 4 | HAE | Forest | Tourist-associated habitat with frequent provisioning and food raiding | Provisioned | High | No |
| H5 | Rhesus macaque | 8 | HAE | Forest | Highway roadside habitat with persistent anthropogenic feeding by road users | Provisioned | High | No |
| H6 | Lion-tailed macaque | 5 | HAE | Fringe | Roadside forest fragment adjacent to settlements with frequent access to anthropogenic foods | Provisioned | Low | No |
| H7 | Bonnet macaque, Rhesus macaque | 12 | HAE | Forest | Temple-associated habitat with provisioning, kitchen raiding and regular human presence | Provisioned | High | Bonnet + Rhesus |
| L1 | Rhesus macaque | 5 | LAE | Forest | Protected forest interior with minimal anthropogenic food exposure | Non-provisioned | Low | No |
| L2 | Rhesus macaque | 11 | LAE | Fringe | Forest-village edge with occasional access to tourist leftovers and seasonal crop resources | Non-provisioned | Low | No |

| | | | | | | | | |
|---|---|---|---|---|---|---|---|---|
| L3 | Bonnet macaque | 5 | LAE | Forest | Protected forest with infrequent and unpredictable anthropogenic food access | Provisioned | Low | No |
| L4 | Lion-tailed macaque | 15 | LAE | Fringe | Plantation-forest mosaic with predominantly natural foraging and minimal provisioning | Non-provisioned | Low | No |
| L5 | Bonnet macaque | 2 | LAE | Rural | Interior temple site visited primarily during religious occasions, resulting in low and unpredictable anthropogenic food access | Provisioned | High | No |

Sampling sites, host species composition, habitat characteristics, provisioning status, predominant human interaction level, and sympatric species occurrence. Sites were classified as high anthropogenic exposure (HAE) or low anthropogenic exposure (LAE) based on the predictability and availability of anthropogenic food resources, provisioning frequency, and observed reliance on anthropogenic foods during sampling. Human interaction level refers to the predominant mode of direct human–macaque interactions at each site. Sympatric sites indicate locations where two macaque species occurred at the same site and were included in sympatric host comparisons.

Table S2. PERMANOVA and multivariate dispersion analyses of gut microbial community structure across anthropogenic environments and sympatric host comparisons.

| Analysis | Comparison | Metric | $R^2$ | F | p-value | Dispersion p-value |
|---|---|---|---|---|---|---|
| Global | Host × environment | Bray–Curtis | 0.024 | 1.53 | 0.0001 | NS |
| Global | Host × environment | Weighted UniFrac | — | — | 0.324 | NS |
| Global | Host × environment | Unweighted UniFrac | 0.025 | 1.59 | 0.0069 | NS |
| Bonnet macaque | HAE vs LAE | Bray–Curtis | 0.027 | — | 0.021 | 0.001 |
| Bonnet macaque | HAE vs LAE | Weighted UniFrac | 0.051 | — | 0.024 | NS |
| Bonnet macaque | HAE vs LAE | Unweighted UniFrac | 0.038 | — | 0.023 | NS |
| Lion-tailed macaque | HAE vs LAE | Bray–Curtis | 0.047 | — | 0.001 | NS |
| Lion-tailed macaque | HAE vs LAE | Weighted UniFrac | 0.229 | — | 0.001 | NS |
| Lion-tailed macaque | HAE vs LAE | Unweighted UniFrac | 0.066 | — | 0.001 | NS |
| Rhesus macaque | HAE vs LAE | Bray–Curtis | 0.051 | — | 0.001 | 0.001 |
| Rhesus macaque | HAE vs LAE | Weighted UniFrac | 0.073 | — | 0.001 | NS |
| Rhesus macaque | HAE vs LAE | Unweighted UniFrac | 0.049 | — | 0.003 | NS |
| H1 sympatry (Bonnet vs LTM) | Host species | Bray–Curtis | 0.049 | 1.61 | 0.0002 | 0.745 |
| H1 sympatry (Bonnet vs LTM) | Host species | Weighted UniFrac | 0.065 | 2.15 | 0.022 | 0.652 |
| H7 sympatry (Bonnet vs Rhesus) | Host species | Bray–Curtis | — | — | 0.367 | 0.033 |
| H7 sympatry (Bonnet vs Rhesus) | Host species | Weighted UniFrac | — | — | 0.583 | NS |

PERMANOVA was used to test differences in microbial community composition among comparison groups. Dispersion p-values were obtained from betadisper analyses and were used to assess the homogeneity of multivariate dispersion.

Figure S2.

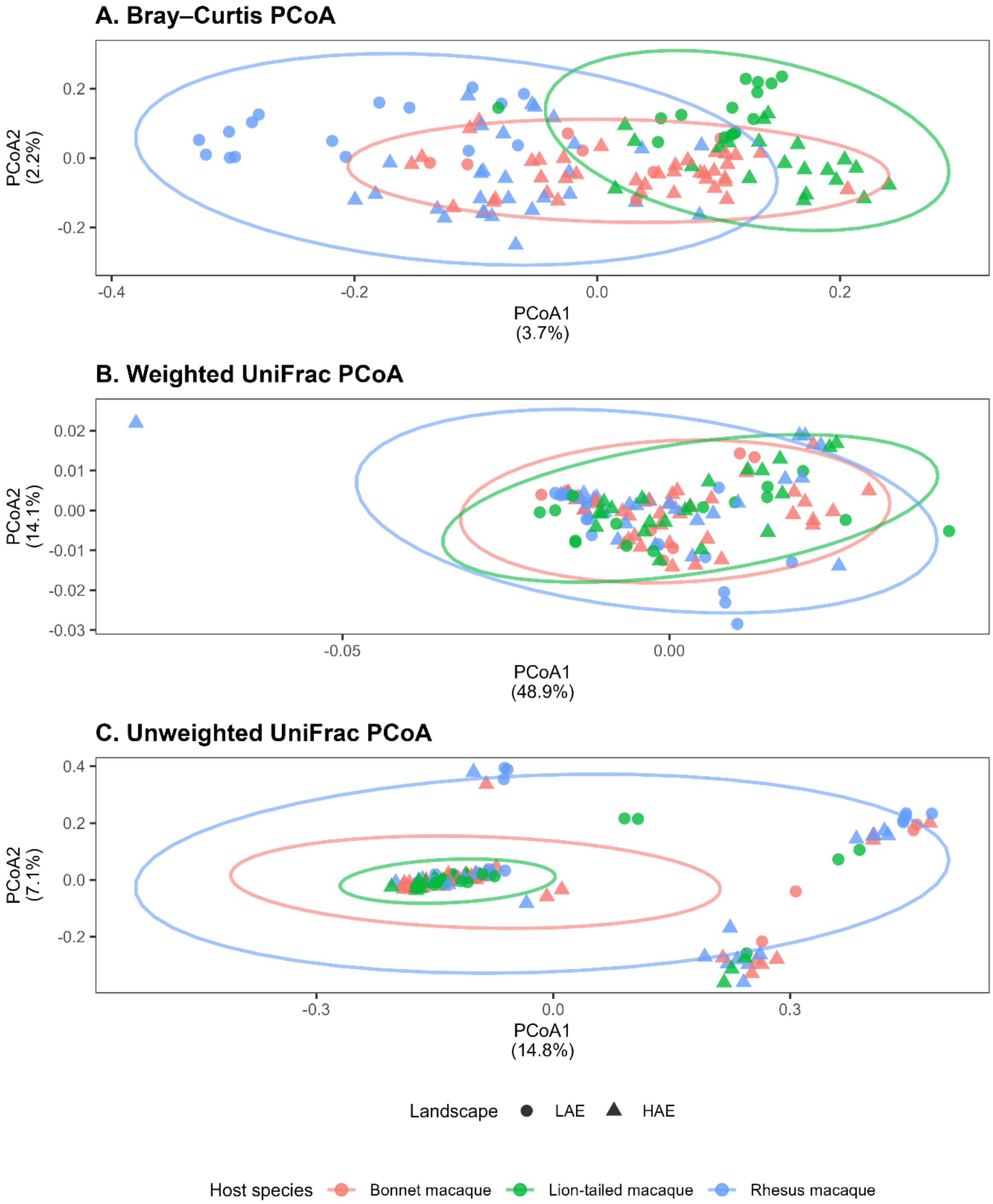


**Figure S2.** Global principal coordinates analysis (PCoA) of gut microbial community composition across all macaque samples based on **(A)** Bray–Curtis dissimilarity, **(B)** weighted UniFrac distance, and **(C)** unweighted UniFrac distance. Colours indicate host species (BM, bonnet macaque; LTM, lion-tailed macaque; RM, rhesus macaque), and symbols indicate habitat category (LAE, low anthropogenic exposure; HAE, high anthropogenic exposure). Ellipses represent the 95% confidence intervals around group centroids for each host species.

Table S3. Differentially abundant amplicon sequence variants (ASVs) associated with anthropogenic environments.

| ASV_ID | Phylum | Family | Genus | LogFC | FDR | Direction |
|---|---|---|---|---|---|---|
| ASV01 | Firmicutes | Ruminococcaceae | Faecalibacterium | 1.168 | 0.000121 | HAE enriched |
| ASV02 | Bacteroidota | Prevotellaceae | Prevotella | 0.731 | 0.00135 | HAE enriched |
| ASV03 | Firmicutes | Lachnospiraceae | [Eubacterium]_hallii_group | -0.913 | 0.00135 | LAE enriched |
| ASV04 | Fibrobacterota | Fibrobacteraceae | Fibrobacter | -0.777 | 0.00194 | LAE enriched |
| ASV05 | Bacteroidota | Prevotellaceae | Prevotellaceae_NK3B31_group | -0.794 | 0.00194 | LAE enriched |
| ASV06 | Bacteroidota | Prevotellaceae | Prevotella | 0.928 | 0.00194 | HAE enriched |
| ASV07 | Firmicutes | Ruminococcaceae | Faecalibacterium | -0.763 | 0.00194 | LAE enriched |
| ASV08 | Firmicutes | Lachnospiraceae | Dorea | -0.68 | 0.00194 | LAE enriched |
| ASV09 | Firmicutes | Peptostreptococcaceae | Intestinibacter | -0.617 | 0.00227 | LAE enriched |
| ASV10 | Firmicutes | Lachnospiraceae | Blautia | -0.677 | 0.0028 | LAE enriched |
| ASV11 | Verrucomicrobiota | WCHB1-41 | WCHB1-41 | 0.663 | 0.00443 | HAE enriched |
| ASV12 | Firmicutes | Lactobacillaceae | Lactobacillus | 0.632 | 0.00443 | HAE enriched |
| ASV13 | Actinobacteriota | Bifidobacteriaceae | Bifidobacterium | -0.62 | 0.00443 | LAE enriched |
| ASV14 | Firmicutes | Lactobacillaceae | Lactobacillus | 0.628 | 0.00709 | HAE enriched |

| | | | | | | |
|---|---|---|---|---|---|---|
| ASV15 | Bacteroidota | Prevotellaceae | Prevotella | 0.599 | 0.00758 | HAE enriched |
| ASV16 | Firmicutes | Butyricicoccaceae | Unclassified | 0.495 | 0.00758 | HAE enriched |
| ASV17 | Bacteroidota | Prevotellaceae | Prevotella | -0.53 | 0.00809 | LAE enriched |
| ASV18 | Firmicutes | Ruminococcaceae | Faecalibacterium | -0.641 | 0.00809 | LAE enriched |
| ASV19 | Firmicutes | Lactobacillaceae | Lactobacillus | 0.49 | 0.00919 | HAE enriched |
| ASV20 | Firmicutes | Ruminococcaceae | Faecalibacterium | 0.489 | 0.0103 | HAE enriched |
| ASV21 | Firmicutes | Lactobacillaceae | Lactobacillus | 0.482 | 0.0155 | HAE enriched |
| ASV22 | Bacteroidota | Prevotellaceae | Prevotella | 0.414 | 0.0202 | HAE enriched |
| ASV23 | WPS-2 | WPS-2 | WPS-2 | 0.482 | 0.0204 | HAE enriched |
| ASV24 | Firmicutes | Lactobacillaceae | Lactobacillus | 0.423 | 0.0273 | HAE enriched |
| ASV25 | Bacteroidota | Prevotellaceae | Prevotella | 0.438 | 0.0273 | HAE enriched |
| ASV26 | Firmicutes | Lactobacillaceae | Lactobacillus | 0.364 | 0.0477 | HAE enriched |
| ASV27 | Firmicutes | Lachnospiraceae | Blautia | -0.386 | 0.0477 | LAE enriched |

Differentially abundant ASVs identified using ANCOM-BC2. Taxonomic assignments are shown at the highest confidently assigned rank. Positive log fold-change values indicate enrichment in high anthropogenic exposure (HAE) environments, whereas negative values indicate enrichment in low anthropogenic exposure (LAE) environments. Statistical significance was assessed using Benjamini–Hochberg false discovery rate correction (FDR < 0.05).

Table S4. Multidimensional microbiome responses to anthropogenic environments across macaque species.

| Microbiome dimension | Bonnet macaque | Lion-tailed macaque | Rhesus macaque |
|---|---|---|---|
| Observed ASV richness | NS | Significant ↓ | NS |
| Shannon diversity | NS | Significant ↓ | NS |
| Faith's phylogenetic diversity | NS | Significant ↓ | NS |
| Bray–Curtis community composition | Significant | Significant | Significant |
| Weighted UniFrac phylogenetic structure | Significant | Significant | Significant |
| Unweighted UniFrac phylogenetic structure | Significant | Significant | Significant |
| Significant differentially abundant ASVs (ANCOM-BC2) | 0 | 1 | 6 |
| Bray–Curtis dispersion heterogeneity | Yes | No | Yes |

Summary of significant responses detected across alpha diversity, beta diversity, phylogenetic structure, differential abundance, and community dispersion analyses. Significance refers to comparisons between low- and high-anthropogenic-exposure environments within each host species. Arrow indicates the direction of alpha diversity change in high anthropogenic exposure (HAE) environments relative to low anthropogenic exposure (LAE) environments.
